# Testing Wright’s Intermediate Population Size Hypothesis – When Genetic Drift is a Good Thing

**DOI:** 10.1101/2022.09.07.506960

**Authors:** Mitchell B. Cruzan

**Affiliations:** Portland State University, Department of Biology

**Keywords:** Adaptative evolution, conservation genetics, drift facilitation, drift load, inbreeding load, population bottleneck, purging, segregation load, simulations

## Abstract

In his 1931 monograph, Sewall Wright predicted genetic drift would overwhelm selection in very small populations, and selection would dominate in large ones, but also concluded drift could facilitate selection in populations of intermediate size. The idea that drift and selection would act together in smaller populations has not been evaluated using analytical or numerical approaches even as empirical evidence of rapid evolution associated with population bottlenecks has continued to accumulate. I used forward-time simulations with random mating and discrete generations to test the hypothesis that drift can facilitate selection in small populations. I find evidence of drift facilitation of selection as increases in levels of *Δq* in small populations (*N*<100) when selection is weak (*s*<0.2) and when allele frequencies are low (*q*<0.5). Fixation of beneficial mutations is accelerated by drift facilitation in small populations for recessive and codominant alleles, and less so for dominant alleles. Drift facilitation accelerated fixation of beneficial mutations in small populations compared to predictions from diffusion equations, while fixation time was longer than predicted in large populations. Drift facilitation increases the probability of fixation of new mutations in small populations. Accumulation of beneficial mutations (fixation flux) over several thousand generations was high in small populations and declined rapidly for large populations, which accumulated large amounts of standing genetic variation. Even though selection is more efficient in large populations, the increased time for allele replacement and lack of drift facilitation can result in substantially slower rates of adaptive evolution. Small populations were more susceptible to the accumulation of drift load, while larger populations maintained higher levels of segregation load. These results indicate that drift facilitation in small populations promotes purging of genetic load and accelerated fixation of beneficial mutations, and may account for the large number of observations of rapid adaptation during population bottlenecks.

Impact Summary – After the recognition of Gregor Mendel’s contributions to our understanding of the inheritance of genetically-determined traits around 1900, there was confusion as to whether the type of variation Mendel studied could account for evolution by natural selection, as described by Charles Darwin. This controversy was resolved when three theoreticians (Ronald Fisher, Sewell Wright, and J.B.S. Haldane) published books that integrated Mendelian genetics with evolution. Their contributions (referred to as the Modern Synthesis), focused on evolutionary processes occurring within and among populations of a species, and established a mathematical foundation for our understanding of evolutionary biology. The mathematical models developed by the three architects of the modern synthesis, and those who followed, predicted that the effects of natural selection would be overwhelmed by random genetic changes (referred to as Genetic Drift) in small populations, and that genetic drift would be minimal, while selection would be most effective in large populations. Even though one of Wright’s major conclusions was that genetic drift and selection would work together (Drift Facilitation) to promote adaptive evolution in intermediate-sized populations, this idea has been almost completely ignored since it was first introduced in 1931. In this study, I use simulations of evolution in natural populations to evaluate the potential for drift facilitation to promote evolution in small populations. My work largely confirms Wright’s predictions; the removal of deleterious mutations and promotion of adaptive evolution are enhanced in population sizes ranging from about 10 to 100. These results indicate that our paradigm for our understanding of evolution within populations needs refinement to emphasize the importance of drift facilitation in small populations, and to recognize that periods of reduced population size are opportunities for enhanced levels of adaptive evolution.

“In a population of intermediate size … there is continual random shifting of gene frequencies … which leads to a relatively rapid, continuing, irreversible, and largely fortuitous, but not degenerative series of changes, even under static conditions.” Wright 1931, Page 157.

## Introduction

The mathematical models for population genetics developed during the Modern Synthesis (Fisher 1930; Wright 1931; Haldane 1932) and beyond (Kimura 1983; Crow 1987; Ewens 2004; Charlesworth and Charlesworth 2017; Charlesworth 2020) have provided a foundation for a general understanding of evolutionary processes accounting for divergence among populations, adaptation, and diversification among lineages. A fundamental aspect of this framework is the dichotomy between the roles of genetic drift and selection; specifically, that drift governs the accumulation of neutral and degenerative variation while only selection can promote the purging of deleterious mutations and the fixation of beneficial mutations (Crow and Kimura 1970; Kimura 1983). The general prediction is that genetic drift will dominate the dynamics of allele frequency change in small populations when selection is weak (i.e. when *s < 1/4N*_*e*_; Charlesworth 2009), resulting in the accumulation of genetic load and limited potential for adaptive evolution. In contrast, selection is expected to overwhelm the effects of drift in large populations leading to fixation of beneficial mutations and the purging of genetic load (Charlesworth 2002; Gossmann et al. 2012; Lanfear et al. 2014; Charlesworth and Charlesworth 2017). However, in his 1931 monograph, Wright emphasized the idea that the rate of fixation of beneficial mutations would be much slower in large populations and that drift could overwhelm selection in very small populations, but there was potential for genetic drift and selection to act together to promote rapid adaptation in populations of “intermediate” size. This idea was intrinsic to his Shifting Balance Theory of adaptive evolution where he predicted that adaptive variants were more likely to become fixed in small populations, and subsequently spread by gene flow (Wright 1932). These predictions were premised on the idea that there was a range of population sizes where both drift and selection are effective enough to allow “fortuitous” shifts in allele frequencies to facilitate selection by generating larger numbers of high-fitness genotypes resulting in accelerated accumulation of adaptive genetic variation. This idea (hereafter referred to as “drift facilitation”) was seized upon by George Gaylord Simpson (1944), who proposed small population size was a likely explanation for the lack of fossilization during rapid transitions that resulted in the origin of novel forms, including new genera and families. This idea of “quantum evolution” (i.e. periods of rapid evolution when selection is facilitated by genetic drift; Simpson 1944) inspired a number of evolutionary biologists, who subsequently identified a wide range of examples of adaptation associated with reduced population size (Mayr 1954; Lewis and Raven 1958). Verne Grant (1963, 1981) expanded on the idea of quantum evolution to include rapid speciation events in small, isolated populations at the periphery of a widespread species’ range (budding or peripatric speciation; Mayr 1954). While interest in the idea of quantum evolution has waned, evidence of rapid evolution associated with population bottlenecks continues to accumulate (e.g., Carvalho et al. 1996; Rosenblum et al. 2007; Tepolt et al. 2009; Marsico et al. 2011; Cruzan 2019; Rego et al. 2019; Chaturvedi et al. 2021; Cruzan et al. 2021; Mahrt et al. 2021; Yin et al. 2021; van der Zee et al. 2022; Appendix 3). Surprisingly, there have been no attempts to assess the potential for drift facilitation of selection in populations of small or intermediate size using analytical or numerical approaches.

In this study, I use forward-time, individual-based simulations of populations to evaluate the potential for drift facilitation to promote evolution in intermediate-sized (small) populations. I evaluate the effects of selection (*s*), dominance (*h*), and population size (*N*) on changes in allele frequencies between generations (*Δq = q*_*n+1*_ *– q*_*n*_) and its consequences for the loss or accumulation of deleterious mutations, and the time to fixation and probability of fixation for beneficial mutations. I specifically focus on short time-frames to account for the numerous observations of rapid evolution within a few hundred generations during population bottlenecks. In all cases, loci are bi-allelic and segregate independently, generations are discrete, populations are dioecious with equal sex ratios, and mating is random (i.e. *N* = *N*_*e*_). I first characterize the effects of population size, selection, and dominance on Δq for beneficial alleles going to fixation, and then examine the consequences of selection and population size for the accumulation of genetic load and beneficial mutations. For genetic load, I examine the proportion of loci that became fixed for deleterious mutations (drift load), and the proportion of deleterious mutations that remain segregating after a number of generations (segregation load). In separate simulations, I examine the probability of fixation for beneficial mutations, the time to fixation, and fixation flux (the rate at which beneficial alleles accumulate; Otto and Whitlock 1997). My results demonstrate the effects of drift facilitation as elevated levels of *Δq* in small populations (*N ≤ 100*) when allele frequencies are low and selection is weak. I find that drift facilitation of selection promotes rapid purging of genetic load and the accumulation of beneficial mutations in small populations while large populations accumulate greater levels inbreeding load and standing genetic variation for beneficial mutations.

## Methods

Forward-time, individual-based simulations were conducted using the SimuPOP module (Peng and Kimmel 2005) in Python and the COEUS High-Performance Computing Cluster at Portland State University. *A* set of *N* individuals, each homozygous for a single locus, were generated using the sim.Population simulator and mutations introduced (*a* → *a*’) that were affected by varying levels of selection (*s*) and dominance (*h*). Transmission of alleles between generations was governed by the sim. MendelianGenoTransmitter. Selection was imposed at the time of mating by defining the fitness of each diploid genotype using MapSelector in SimuPOP (all Python scripts are available in Appendix 1). Depending on the goals of the simulation, mutations were either introduced once in the first generation, or at a per-generation rate determined by *µ*. Each population size of *N* individuals was replicated *k* times. For each replication, the proportion of individuals with fixed mutations (*q*_*fixed*_), and the average allele frequency across individuals 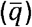 were extracted after g generations. Two sets of simulations were run; the first set examined levels of drift and segregation load, and the second set focused on beneficial mutations by evaluating the proportion of loci fixed for mutations, the rate of fixation of individual mutations, and the fixation flux over g generations.

### Testing for Drift Facilitation of Selection in Small Populations

I tested for the effects of drift facilitation of selection by comparing *Δq* (*Δq* = *q*_*n+1*_ *– q*_*n*_) in small (*N = 50*) and large (*N = 1000*) populations across generations as beneficial mutations increased from their initial frequency of *q = 1/2N* to fixation. Mutations were repeatedly generated and their allele frequencies tracked until they were fixed or lost (Python script available in Appendix 1.A). For mutations that fixed, *Δq* for each of the n generations was calculated along with its mean and standard deviation across generations. Mean *Δq* for mutations that fixed was compared for a range of selection coefficients (*s = 0*.*01, 0*.*04, 0*.*12*, and *0*.*20*) across population sizes ranging from *N = 4* to *N = 3000* for recessive, additive, and dominant mutations (*h = 0*.*0, 0*.*5*, and *1*.*0*). I evaluated the effects of the effects of the strength of selection (*s = 0*.*01* to *0*.*20*) in a small (*N = 100*) and large (*N = 1000*) population on the mean and standard deviation of *Δq (k = 10,000* replications) for beneficial mutations going to fixation (Appendix 1B).

### Consequences of Drift Facilitation for Genetic Load

The frequency of fixed deleterious mutations (drift load) across loci and of the proportion of loci segregating for deleterious mutations after g generations (segregation load) was assessed for population sizes ranging from *N* = 2 to 5000 (Python script available in Appendix 1C). For each of *k = 1000* replications for each population size, s was randomly generated from a Poisson distribution (range: 0.0 – 0.25; 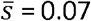), and h was generated from a uniform distribution between 0.0 and 0.2 (assuming that deleterious mutations are generally recessive with 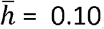; Willis 1999; García-Dorado and Caballero 2000; Agrawal and Whitlock 2011; Ruzicka et al. 2021). Fitness of the *aa, aa’*, and *a’a’* genotypes were set to 1.0, 1-*hs*, and *1-s*, respectively. The initial allele frequency of *a’* was set at *q* = 0 and mutations were allowed to accumulate for *g = 1000* generations with *µ = 1×10*^*-3*^ This assumes that each locus consists of thousands of base pairs (i.e. if the per bp/generation mutation rate is on the order of *1×10*^*-9*^), but the same qualitative results are obtained for lower mutation rates over longer timeframes.

The proportion of loci fixed for deleterious mutations (*q*_*fixed*_) and selection (s) for each mutation were used to estimate the fitness effects of fixed deleterious mutations (drift load) for each replication of each population size as the reduction in fitness due to the selection coefficient weighted by the proportion of loci fixed for the mutation;

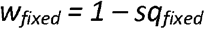

Segregation load was assessed as the inbreeding effects on fitness for segregating mutations (inbreeding depression; Charlesworth and Willis 2009; Willi et al. 2013) for 1000 replicates for each population size after 1000 generations. Inbreeding depression was estimated based on the mean fitness of hypothetical progeny that would be the result of self-fertilization compared to progeny from random outcrossing. Fitness after selfing was estimated as,

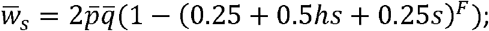

where 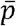 and 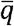 are the mean frequencies of the normal (*a*) and mutant (*a’*) alleles across segregating loci (individuals), and *F* is the number of segregating sites at the end of 1000 generations. I calculated the expected fitness of progeny from random outcrossing within each population as,

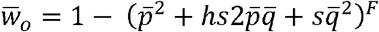

Segregation load was then estimated for each *N* as inbreeding depression for selfed progeny compared to outcrossed progeny (Charlesworth and Willis 2009) as,

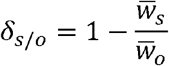

These estimates of drift load and segregation load assume independent fitness effects across unlinked loci.

### Consequences of Drift Facilitation for Adaptation

I examined the processes of adaptation in populations using simulations to evaluate the probability of fixation of beneficial mutations, the rate of fixation of individual alleles, and the fixation flux for a range of population sizes (*N*), selection (*s*), and dominance (*h*; Appendix 1D). For beneficial mutations (*a* → *a’*), the fitness of the *aa, aa’*, and *a’a’* genotypes were set to *1-s, 1-hs*, and *1*.*0*, respectively.

The first set of simulations compared the fixation of beneficial alleles across a range of initial allele frequencies from *q = 0*.*01* to *0*.*50* when *Ns = 2* for population sizes of *N = 100 (s = 0*.*02)* and *N = 1000 (s = 0*.*002)*. Initial frequency of the a’ allele was set at *q = 1/2N (µ = 0)* and the proportion of loci fixing for the mutant allele was compared to the diffusion prediction for the probability of fixation 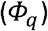 derived by Kimura (1957; 1962). Alleles were codominant in simulations (*h = 0*.*5*) to match the assumptions of the diffusion approximation;

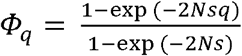

A second set of simulations examined the number of generations required for fixation over population sizes ranging from *N = 4* to *3000* with s randomly drawn from a Poisson distribution (*s = 0*.*0 to 0*.*25*) and h drawn from a uniform distribution (*h = 0*.*0* to *1*.*0*). Initial frequency of the *a’* allele was set at *q = 1/2N (µ = 0)* and the proportion of loci fixing for the mutant allele was recorded after 10,000 generations. I compared the number of generations required for fixation for beneficial alleles (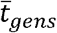 across *k = 10* replicates per population size) to the predicted time to fixation 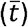 from the diffusion approximation (Ewens 1979; Otto and Whitlock 2013);

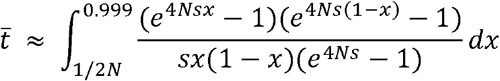

This integral is undefined when the upper limit is 1.0, and the approach to fixation is asymptotic, so an arbitrary upper limit of 0.999 was chosen for comparison to fixation frequencies from the simulations.

While this approach describes fixation times for individual alleles, it does not account for the contribution of the number of mutations entering populations each generation based on population size for the accumulation of beneficial mutations over time (fixation flux; Otto and Whitlock 1997). We expect flux to be affected by the probability of fixation for individual mutations and the number of mutations entering the population each generation. I evaluated the probability of fixation across a range of population sizes (*N = 4* to *3000*) by introducing random mutations (*s = 0*.*0* to *0*.*2* following a Poisson distribution, and *h = 0*.*0* to *1*.*0* following a flat distribution) to 1000 loci in the first generation and tracking the proportion that fix after 10,000 generations (100 replications per population size).

We expect that more mutations would occur in larger populations and consequently flux should increase with population size. On the other hand, fixation will take longer in large populations because of the time required for replacement of the original allele in all individuals. Since the goal of this study is to understand conditions leading to rapid adaptation to a novel environment, simulations were limited to a few thousand generations without a prior burn-in period to allow for the accumulation of standing genetic variation. I examined fixation flux for population sizes ranging from *N = 4* to *3000* with *µ = 1×10*^*-4*^, *Ns = 2 (s = 2/N)*, and *h = 0*.*5* with 1000 loci and 10 replicates per population size for time periods of *g =1000, 2000*, and *4000* generations (Appendix 1E). Fixation flux under selection was compared to fixation with only genetic drift (*s = 0*) over 1000 generations across the same range of population sizes. In addition, I estimated the mean and standard deviation of Δq across generations for beneficial mutations that ultimately fixed in populations across a range of populations sizes (*N = 4* to *3000*) and selection coefficients (*s = 0*.*01* to *0*.*20* for *N = 100* and *1000*).

## Results

*Drift Facilitation of Selection in Small Populations* – Drift facilitation was evident as elevated levels of Δq when population size was small (*N = 50*) compared to large populations (*N = 1000*; Fig. 1). The effects of drift facilitation were strongest when selection was weak (*s* < 0.2), mutations were recessive (h = 0.0), and allele frequencies were low, but there were strong effects of drift facilitation for codominant mutations (Fig. 1), and weaker effects when mutations were completely dominant (Fig. S1). There was a general trend towards the strongest effects of drift facilitation in small populations with weaker selection and lower dominance. Mean *Δq* increased with the strength of selection (Fig. S2) and remained consistently higher in small (*N = 100*) compared to large (*N = 1000*) populations. The standard deviation of *Δq* was consistently higher for small populations and remained fairly constant across a range from *s = 0*.*01* to *0*.*20* (Fig. S3).

**Fig. 1.**
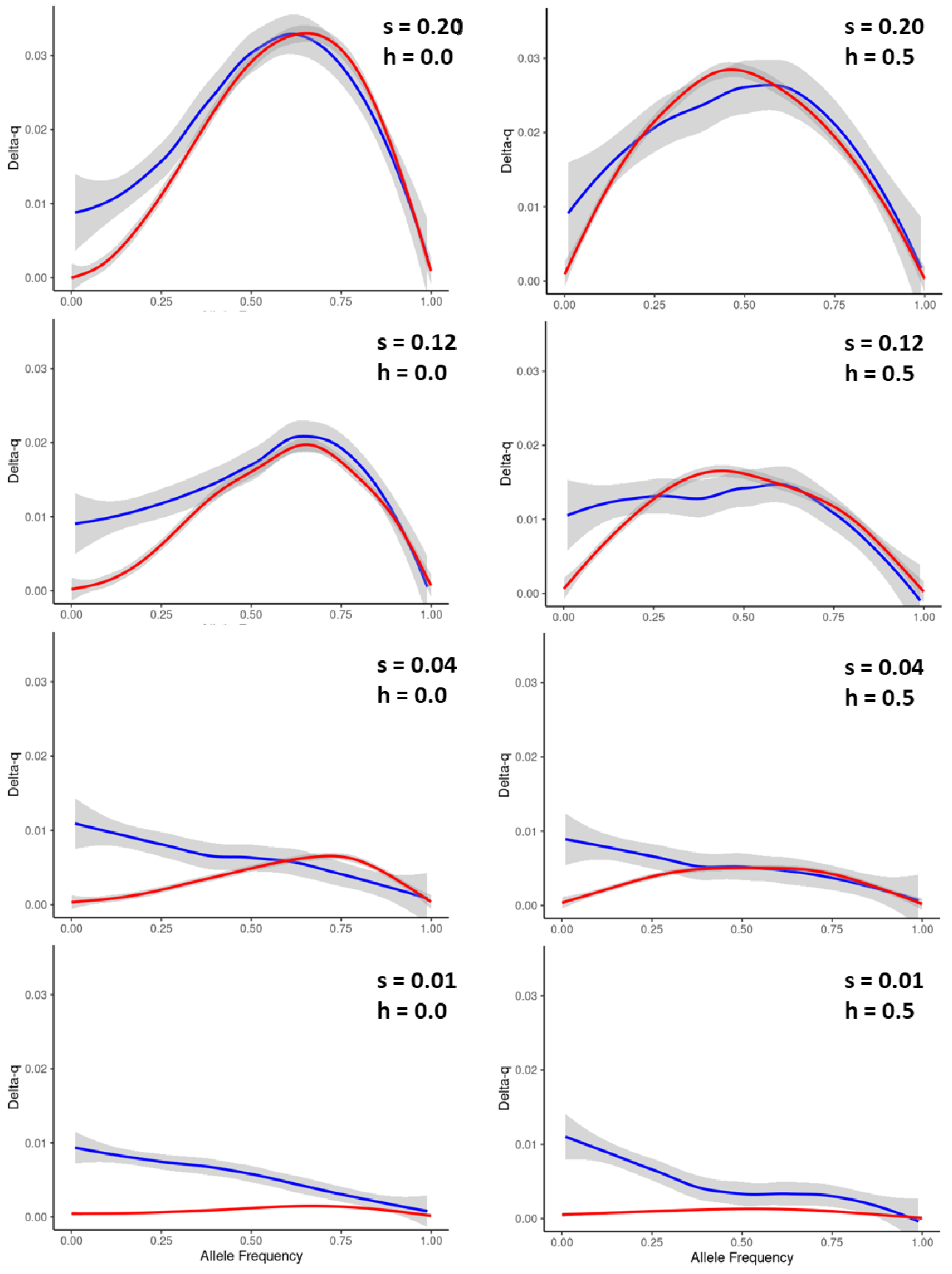
Mean *Δq* across allele frequencies from *q = 1/2N* to *q = 1*.*0* for beneficial alleles in a small (*N = 50*; blue line) compared to a large (*N = 1000*) population for recessive (*h = 0*.*0*; left column) and codominant (*h = 0*.*5*; right column) mutations with selection ranging from *s = 0*.*01* to *s = 0*.*20*.

### Consequences for Genetic Load

Fixation frequencies of deleterious mutations were highest for small populations (< 100), and was effectively zero for population sizes larger than *N = 3000* (Fig. 2A). Mutations fixing in populations when *N > 100* had negligible fitness effects so that drift load was near zero when population size was greater than *N = 100* (Fig. 2B). Larger proportions of loci continued to segregate for deleterious mutations after 1000 generations in populations of less than *N = 10* and more than *N = 100*, while segregation load was lower when *N > 10* and *N < 100*; Fig. 3).

**Fig. 2.**
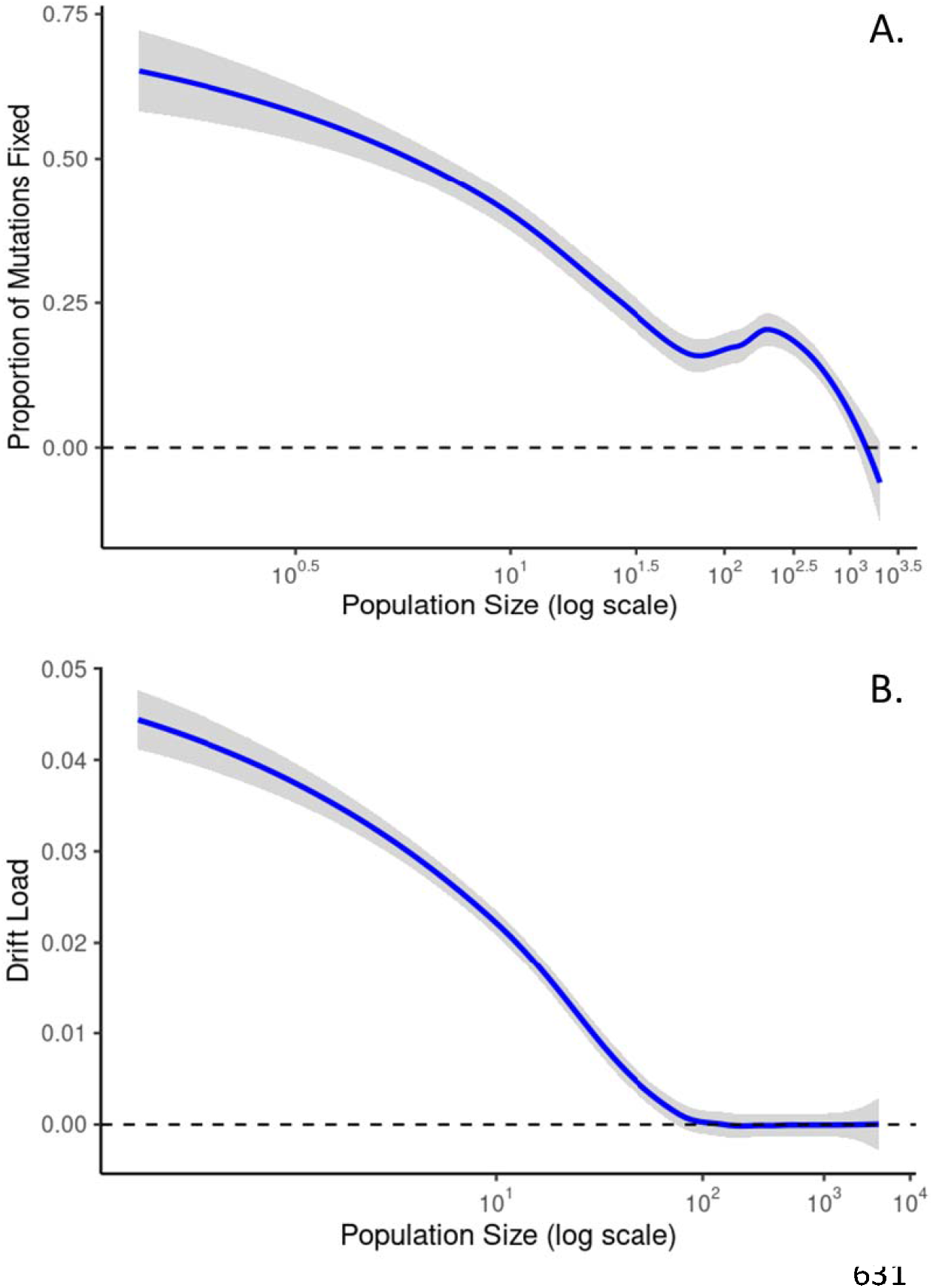
The proportion of deleterious mutations fixing (A) and the levels of drift load (B) across population sizes of *N = 2* to 4000 after 1000 generations (*µ = 1×10-3*). Selection on mutations ranged from 0.0 to 0.25 and were drawn randomly from a Poisson distribution, and dominance ranged from 0.0 to 0.2 and were drawn randomly from a uniform distribution. The blue lines represent the loess-smoothed means from 1000 replications for each *N*, and gray shading indicate 95% confidence intervals.

**Fig. 3.**
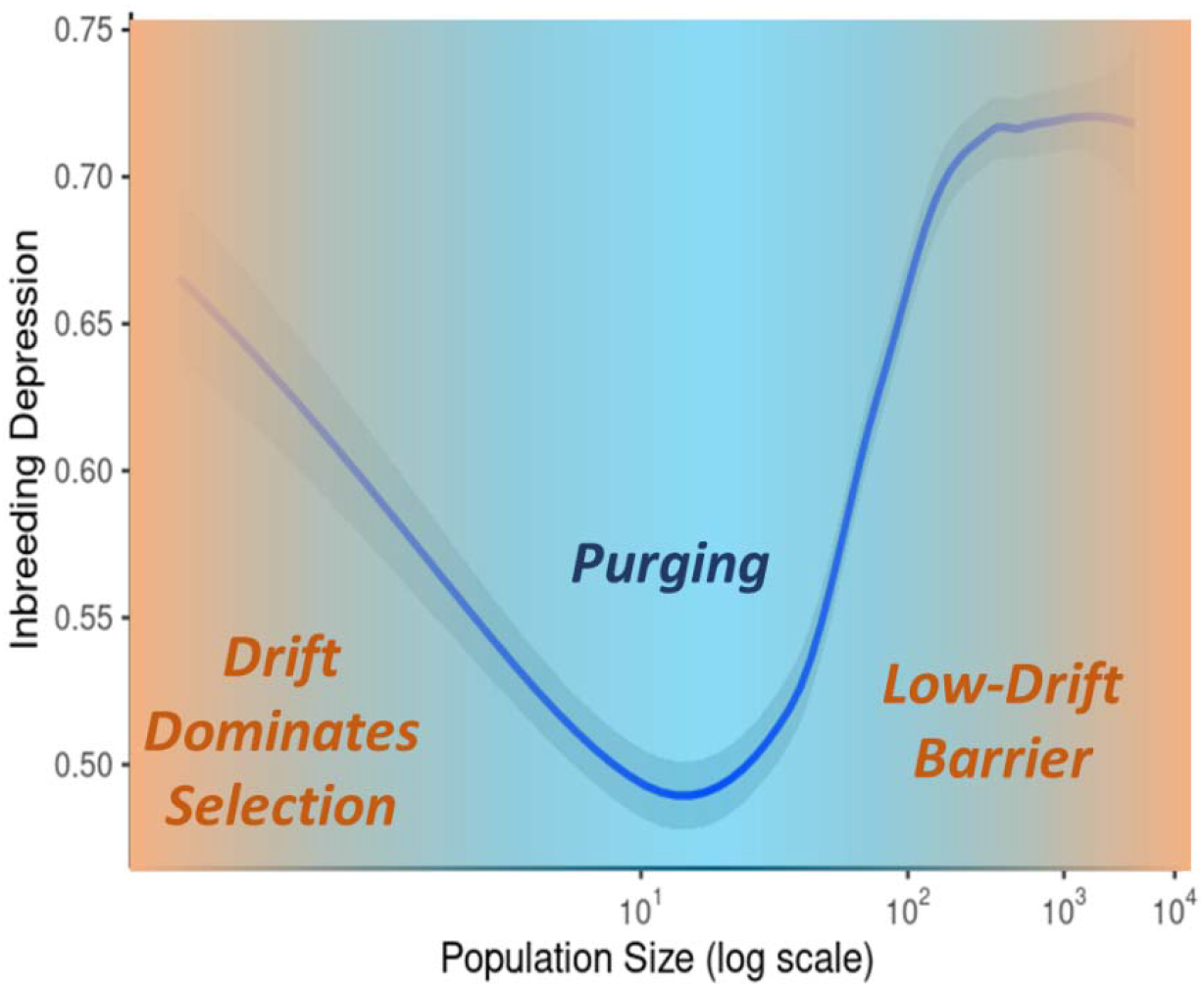
Levels of inbreeding load due to loci segregating for deleterious mutations across population sizes of *N = 2* to 4000 after 1000 generations (*µ = 1×10*). Selection on mutations ranged from 0.0 to 0.25 and were drawn randomly from a Poisson distribution, and dominance ranged from 0.0 to 0.2 and were drawn randomly from a uniform distribution. The blue line represents the loess-smoothed means from 1000 replications for each *N*, and gray shading indicate 95% confidence intervals.

### Consequences for Adaptive Evolution

There was a close fit between the diffusion approximation for the probability of fixation across initial allele frequencies *(Фq)*and the proportion of loci fixed for beneficial mutations with population sizes of 100 and 1000 with *Ns = 2* and codominance (*h = 0*.*5*; Fig. S4).

The time to fixation of individual alleles 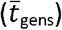 was similar to the diffusion approximation 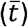 when selection was stronger (*s = 0*.*1* and *0*.*2*) and population size was less than *N = 100*, but was slower than predicted when *N > 100*. Time to fixation was substantially faster when *s = 0*.*04* and *0*.*02* when population sizes were greater than *N = 10* and less than *N = 100* (Fig. 4). The probability of fixation for beneficial mutations was highest for small populations (*N < 100*), and remained lower for large populations (Fig. 5). Fixation of random beneficial mutations was positively associated with the strength of selection (*t = 18*.*79; P < 0*.*0001*) and negatively associated with dominance (*t = -8*.*87; P < 0*.*0001*) and population size (*t = -2*.*71; P = 0*.*0072*). Fixation flux for beneficial mutations with *Ns = 2* was greater after 4000 generations compared to 2000, and 1000 generations (Fig. 6A). Fixation flux and the loss of beneficial mutations was high when *N < 100*, but generally declined with increasing population size (*h = 0*.*5*; Figs. 6A, B) due to the limited time-frame (i.e. < 4000 generations) and the number of generations required for allele replacement in large populations. The number of mutations that remained segregating in large populations was not strongly affected by the length of the time for the accumulation of mutations, and was consistently low when *N < 100*, and increased rapidly when *N > 100* (Fig. 6C).

**Fig. 4.**
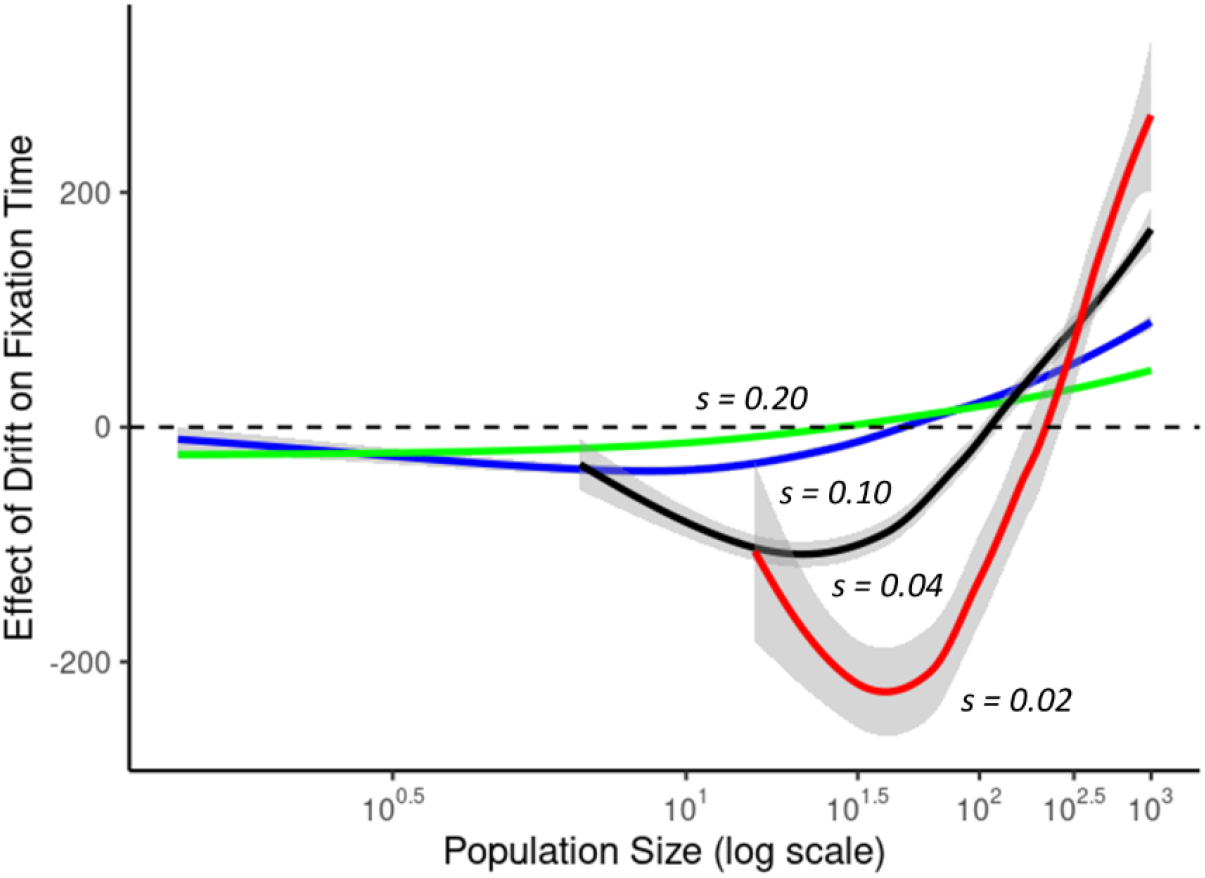
Effects of population size on number of generations for fixation of beneficial alleles from simulations (*h = 0*.*5*) compared to the diffusion approximation averaged across 10 replicates per population size (–) for *s = 0*.*02* (red), *0*.*04* (black), *0*.*10* (blue), and *0*.*20* (green). The lines represent the loess-smoothed means across 100 replications for each *N*, and gray shading indicates 95% confidence intervals.

**Fig. 5.**
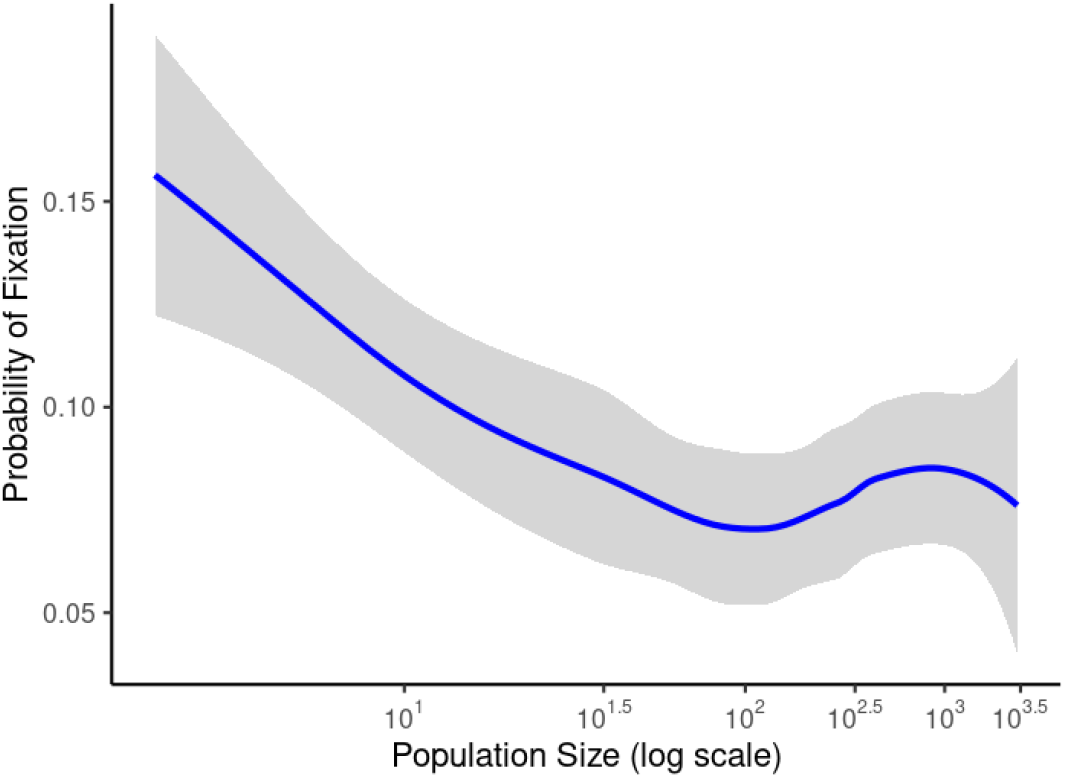
The proportion of loci fixing for random beneficial mutations introduced in the first generation with *s = 0*.*0* to *0*.*2* (Poisson distribution) and *h = 0*.*0* to *1*.*0* (flat distribution) after 10,000 generations for population sizes ranging from *N = 4* to *3000*.

**Fig. 6.**
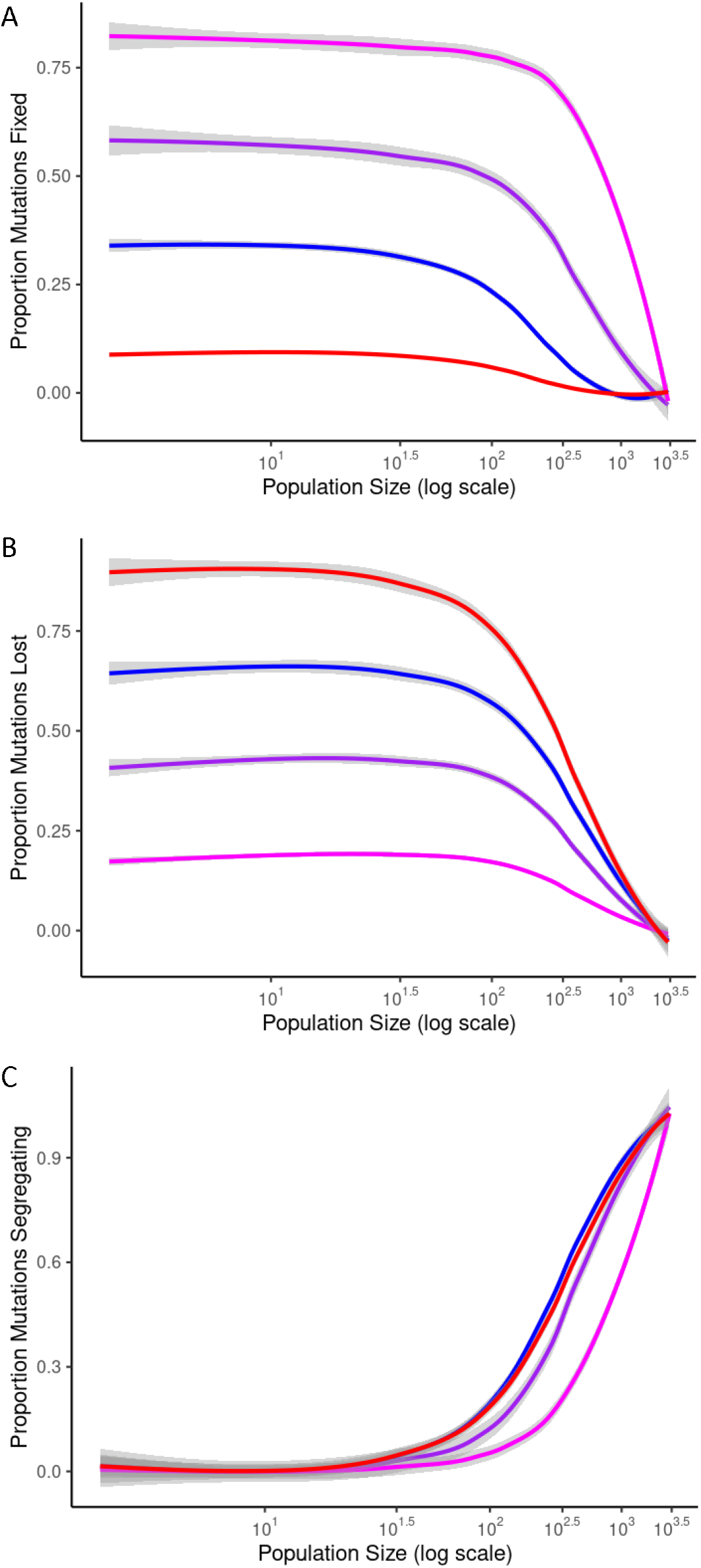
Accumulation of beneficial mutations (A; fixation flux), mutations lost (B), and the proportion continuing to segregate (C) after 1000 (blue), 2000, (purple), and 4000 (magenta) generations with *Ns = 2*, and over 1000 generations with *s = 0* (red) with no prior burn-in. The lines represent the loess-smoothed means across 100 replications for each *N*, and gray shading indicates 95% confidence intervals.

## Discussion

Forward-time simulations of natural populations indicate that drift facilitation of selection elevates *Δq* in small populations (*N ≤ 100*) resulting in lower segregation load, accelerated times to fixation, and higher probability of fixation for beneficial mutations. Drift load was relatively high for small populations, but was negligible for populations larger than *N = 100*, while segregation load was lowest for population sizes between 10 and 100. Results of simulations for beneficial mutations were consistent with diffusion approximations of the probability of fixation when *h = 0*.*5* (Kimura 1957; Kimura 1962). Observed times to fixation matched the diffusion approximation 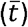 fairly well when selection was strong (*s ≥ 0*.*10* and *h = 0*.*5*), but fixation times were substantially faster when *s < 0*.*10* for population sizes between *N = 10* and 100, and slower for population sizes of *N > 100*. Drift facilitation in smaller populations led to higher probabilities of fixation and high levels of fixation flux of beneficial mutations across a broad range of small population sizes while flux rapidly declined for populations larger then *N = 100* after 1000 generations, and *N = 300* after 4000 generations, which was apparently due to the extended time to fixation in larger populations. While we expect that more beneficial mutations will occur in large populations, the lack of genetic drift reduces the chances of fixation and increases the time to fixation compared to small populations. These results suggest the that drift facilitation provides some explanation for observations of rapid adaptation during periods of reduced population size by increasing purging of deleterious alleles and promoting the acquisition of adaptive genetic variation.

The results presented here generally support Wright’s (1931) predictions about the potential for interactions between drift and selection to promote rapid fixation of beneficial alleles, but the population sizes where drift facilitation of selection is effective are smaller than he predicted. Wright specifically defined intermediate populations as ranging between *Ns = 2*.*5* and *20* (page 148). Simpson (1944) also assumed that the dynamics described by Wright would apply to very large population sizes (i.e. up to N = 25,000) compared to the range of population sizes where drift facilitation was found to be effective in the simulations presented here. In contrast to the predictions of Wright and Simpson, higher rates and probabilities of fixation of beneficial mutations were observed in simulations for *N < 100*. The results presented here indicate that we cannot assume there is a linear tradeoff between the effects of population size and the strength of selection when population sizes small (i.e. for *N < 100*). This is an important result because Ns has been used to represent the compensatory effects between genetic drift and selection in analytical models including diffusion equation analyses of population genetic processes (Kimura 1957; Kimura 1962; Ewens 2004). Given the results presented above, more careful evaluation of the separate effects of selection and population size may be warranted when small populations are being considered.

While population genetic models have mostly ignored the potential for selection in small populations, empirical evidence for the effectiveness of selection during population bottlenecks continues to accumulate. For example, a search of Web of Science using the keywords “rapid and (evolution or adaptation) and bottleneck” yielded 153 publications since 1992 on a wide variety of organisms after removing reviews, editorials, non-biological studies, and studies focused on cancer (details in Appendix 3). A sample of the reported cases of rapid adaptation to novel conditions during establishment and bottlenecks includes artificial introductions of Trinidadian guppies (Carvalho et al. 1996; van der Zee et al. 2022), Eastern fence lizards on novel substrates in New Mexico (Rosenblum et al. 2007), European green crabs on the west coast of North America (Tepolt et al. 2009), the cactus-feeding moth, *Cactoblastis cactorum*, in Florida (Marsico et al. 2011), steelhead trout in Lake Michigan (Willoughby et al. 2018), the invasive grass, *Brachypodium sylvaticum*, in Oregon (Cruzan 2019; Marchini et al. 2019), during evolutionary rescue in the seed beetle *Callosobruchus maculatus* (Rego et al. 2019), in founding populations of *Daphnia* (Chaturvedi et al. 2021), and after migration into Lake Champlain by the Atlantic sea lamprey (Yin et al. 2021). In one case, there was rapid evolution of antibiotic resistance in *Pseudomonas aeruginosa* even when population bottlenecks were severe and selection was weak (Mahrt et al. 2021). Rapid acquisition of adaptive genetic variants also seems to have occurred in a case of recent budding speciation (peripatric speciation; Mayr 1954) in a range-restricted species of buttercup (Cruzan et al. 2021). In all of these cases, adaptation to novel conditions appears to have occurred quickly during periods of reduced population size and suggests drift facilitation of selection may be a widespread phenomenon.

One of the more surprising results of simulations was the observation of higher rates of fixation flux of beneficial mutations when *N < 300*, suggesting drift facilitation can enhance rates of adaptation in small populations. This result is in consistent with diffusion analyses, which predicts flux estimated as *4NsU* (where *U* is the genome-wide mutation rate) will remain constant across population sizes if *Ns* is held constant (Otto and Whitlock 1997; Ewens 2004), but the decline in flux when *N > 300* is due to the slow accumulation of standing genetic variation. The simulations conducted here assumed transitions to novel conditions or colonization events so adaptation from novel mutations would only be for those occurring after the transition. The rate of flux may have been equally high for large populations if substantial “burn-in” periods (> 10,000 generations) had been implemented to allow the accumulation of standing genetic variation. However, the rate of adaptation from standing genetic variation in large populations will depend on initial allele frequency, and even for alleles that are at mid-frequencies (e.g., Lai et al. 2019), the time to fixation would be substantially longer than for small populations.

The observations of elevated *Δq*, reduced time to fixation, and increased probability of fixation of beneficial mutations in small populations are consistent with some previous analyses that predicted genetic drift and selection could interact to increase rates of divergence among populations (Cohan 1984; Lynch 1986), and with selection experiments in maize and simulations where enhanced rates of fixation of small effect mutations were observed when subjected to high genetic drift (Desbiez-Piat et al. 2021). It’s notable that predictions from diffusion analyses are often evaluated only for a range of population sizes that are much larger than what was considered here (e.g., for N > 1000; Charlesworth 2020), but many species have historically experienced severe population bottlenecks. While diffusion analyses have led to numerous insights for evolution in large populations, alternative approaches may be required when population sizes less than *N = 300* are considered.

Previous studies that have found evidence of drift facilitation did not identify its causes (e.g., Cohan 1984; Lynch 1986), but the evaluation of *Δq* across allele frequencies and population sizes sheds some light on the mechanisms responsible for enhancement of reduced times to fixation in small populations. The mean and standard deviation of *Δq* increased in small populations when selection was weak (*s < 0*.*2*) and mostly for allele frequencies of *q < 0*.*5*. For mutations at low frequencies, it appears that “fortuitous” fluctuations due to drift generate larger numbers of high-fitness genotypes (i.e. homozygotes when *h = 0*, and heterozygotes as well when *h = 0*.*5*). Consequently, the effects of drift facilitation are greatest for recessive and codominant mutations as drift facilitation elevates *Δq* and generates a “ratcheting effect” that accelerates changes in allele frequencies. This idea is supported by the observation that the mean of *Δq* is positively associated with selection, and is consistently higher in small compared to large populations. The effects of drift facilitation are apparently different when allele frequencies are close to *q = 0* (for deleterious mutations) or fixation (i.e. *q = 1* for beneficial mutations). Previous analyses indicate that diffusion equations do not perform well when allele frequencies are close to fixation when alleles under selection are completely recessive or dominant (Charlesworth 2020). Simulations performed here indicate that higher levels of drift in smaller populations increase the probability for random fluctuations to result in absorption, while minimal drift in larger populations results in persistent segregation. Consequently, larger populations appear to accumulate higher levels of segregation load and standing variation for beneficial mutations, while having lower drift load lower probabilities of fixation for beneficial mutations compared to smaller populations (i.e. for *N < 100*).

The observation that large populations maintain higher levels of segregation load and standing variation for beneficial mutations suggests that episodes of reduced population size may be disproportionally important for rapid evolution in novel or changing environments. While it remains true that more beneficial mutations are likely to occur in large populations, the number of generations required for fixation may be too great to keep pace with current rates of climate change. On the other hand, large populations possess higher levels of standing genetic variation so bottlenecks would have enhanced potential for the fixation of beneficial mutations (e.g., Chaturvedi et al. 2021). Moreover, reductions in population size due to bottlenecks and founder events are likely to be associated with increased environmental stress, and consequently, we expect selection to often increase during population bottlenecks (Fowler and Whitlock 2002; Matute 2013). Higher frequencies of segregating mutations along with drift facilitation may be responsible for elevated levels of purging following population bottlenecks (Bouzat 2010), and may help explain the substantial number of cases of rapid adaption during episodes of reduced population size (e.g., Carvalho et al. 1996; Rosenblum et al. 2007; Tepolt et al. 2009; Marsico et al. 2011; Cruzan 2019; Rego et al. 2019; Cruzan et al. 2021; Mahrt et al. 2021; Yin et al. 2021; van der Zee et al. 2022; Appendix 3). On the other hand, we expect population bottlenecks and founder events to result in elevated accumulation of drift load (Lynch et al. 1995; Willi et al. 2013), but the simulations presented here indicate the potential for purging of segregation load and the fixation of deleterious mutations depends on the severity of the reduction in population size. It is notable that the standard guideline of a minimal effective population size of N = 50 (i.e. the 50/500 rule; Jamieson and Allendorf 2012) for conservation efforts is too small to prevent the accumulation of drift load, but is in the right range for drift facilitation of beneficial mutations. Further analyses may help resolve more specific effects of changes in population size for adaptation to novel conditions and for the purging of deleterious mutations.

## Conclusions

The results of individual-based, forward-time simulations are consistent with a wide range of observations from natural populations, but contrast with many of the predictions from models based on diffusion equations (e.g., Ewens 2004) when population sizes are small. In general, it appears that Wright’s (1931) intuitive predictions about the potential for genetic drift to facilitate selection were correct, but the population sizes where drift facilitation is most effective are smaller than he suggested. Furthermore, while Wright saw the potential for weak deleterious mutations to fix in small populations, he did not anticipate that drift facilitation could lead to more effective purging of segregation load. One of the most striking results from these simulations is that the probability of fixation and flux of beneficial mutations are substantially high in small populations (*N < 100*) and is similar to flux in larger populations under the assumption that *Ns* remains constant. The assumption that selection is stronger in small populations may generally hold true; especially under conditions where population declines are precipitated by increased levels of environmental stress. Moreover, the time to fixation for beneficial mutations was lower in small populations when selection was weak (*s < 0*.*2*), so the prediction of rapid evolution due to drift facilitation is not dependent on the assumption of increased selection in small populations.

The potential for rapid evolution in small populations may be further enhanced as large populations experience reductions in population size. While it is generally assumed population bottlenecks will result in decreased fitness due to the accumulation of genetic load, and that there will be limited potential for adaptive evolution, this is not always true (Bouzat 2010). The simulation results presented here indicate drift facilitation of selection in small populations leads to effective removal of segregating deleterious mutations, and increases the rate and probability of fixation for beneficial mutations. While there has been some speculation and evidence for enhanced evolution in small populations (Simpson 1944; Grant 1981; Cohan 1984; Lynch 1986), no previous study has provided a detailed characterization of the potential for drift facilitation to promote rapid evolution.

Results from these simulations emphasize the importance of the adaptive evolution potential in small populations. Furthermore, more attention should be paid to the consequences of deleterious mutations as they depend heavily on population size; a clear distinction needs to be made between the fixation of deleterious mutations contributing to drift load in small populations, and the persistence of deleterious mutations in large populations resulting in inbreeding depression due to segregation load. The results of these simulations highlight genetic drift as a generative, beneficial process that promotes evolution in small populations. Alternatively, the lack of drift in large populations effectively stalls purging of segregation load and the fixation of beneficial mutations. While further analyses of drift facilitation are needed, the results presented here give more weight to Wright’s (1931, 1932) suggestion that adaptive evolution is more likely to occur in small populations (Wade and Goodnight 1998), then to Fisher’s (1930) argument that large populations are more important for fixation of beneficial mutations (Coyne et al. 1997). Further work on evolution in small populations from modeling and empirical perspectives will clarify the potential for drift facilitation to promote evolution, and ultimately determine whether the current paradigm for our understanding of population genetic processes of evolution needs to be refined.

## Supporting information

Scripts

Literature search results

## Acknowledgements

I thank Michael Wade and Michael Whitlock for suggestions on the development of simulations and relevant parameters to focus on. Karla de Lima Berg, Elizabeth Hendrickson, Jaime Schwoch, and Ariana Walczyk provided suggestions to improve the manuscript and presentation of results. Thanks to Bo Peng for assistance with coding in simuPop, and to Jim Stapleton and the PSU OIT for assistance with configuration of Coeus HPC for simulations in simuPop. This research was supported by NSF-DEB award 2051235.

## Author Contributions

MBC is solely responsible for the development of the SimuPOP simulations in Python, data analyses, and writing of the manuscript.

## Data Accessibility

All Python scripts are available in Appendix 1 and will be made available on Github.

## Appendix 2. Supplemental Figures

**Fig. S1.**
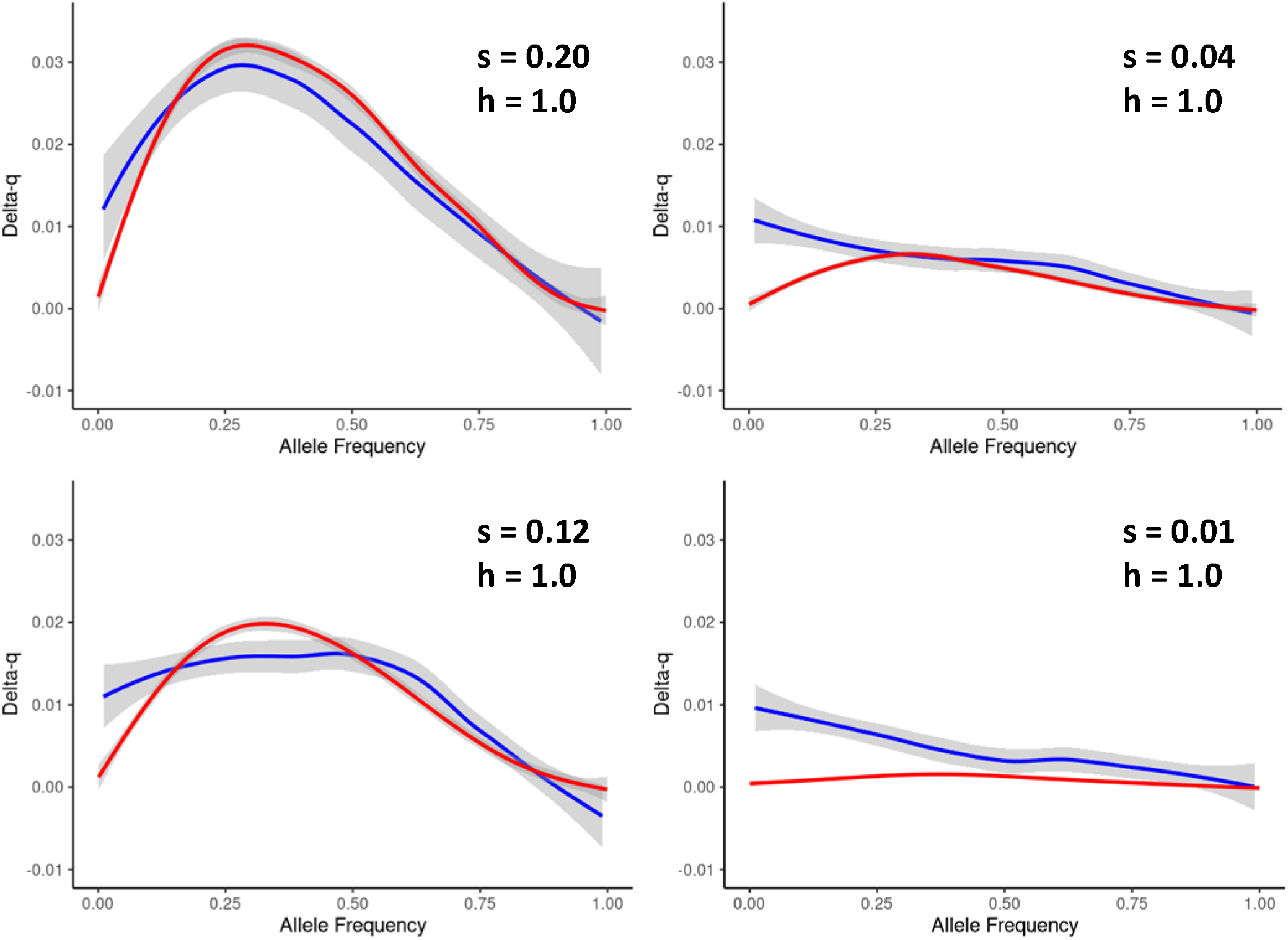
Mean *Δq* across allele frequencies from *q = 1/2N* to *q = 1*.*0* for beneficial alleles in a small (*N = 50*; blue line) compared to a large (*N = 1000*; red line) population for dominant mutations (*h = 1*.*0*) with selection ranging from *s = 0*.*01* to *s = 0*.*20*.

**Fig. S2.**
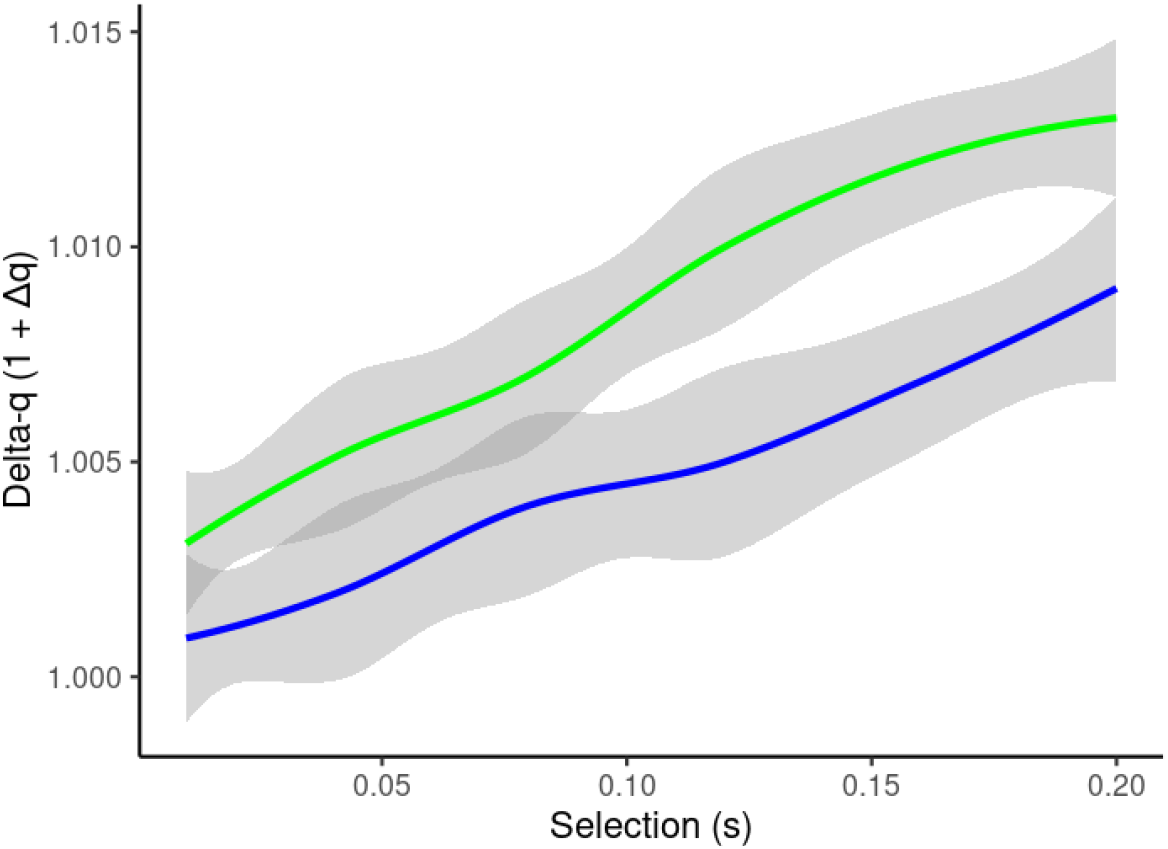
Mean *Δq* for beneficial mutations going to fixation from an initial frequency of *1/2N* across selection coefficients from *s = 0*.*01* to *0*.*20* for *N = 100* (green line) and *N = 1000* (blue line). The line represents the loess-smoothed means across 100 replications for each *N*, and gray shading indicates the 95% confidence interval.

**Fig. S3.**
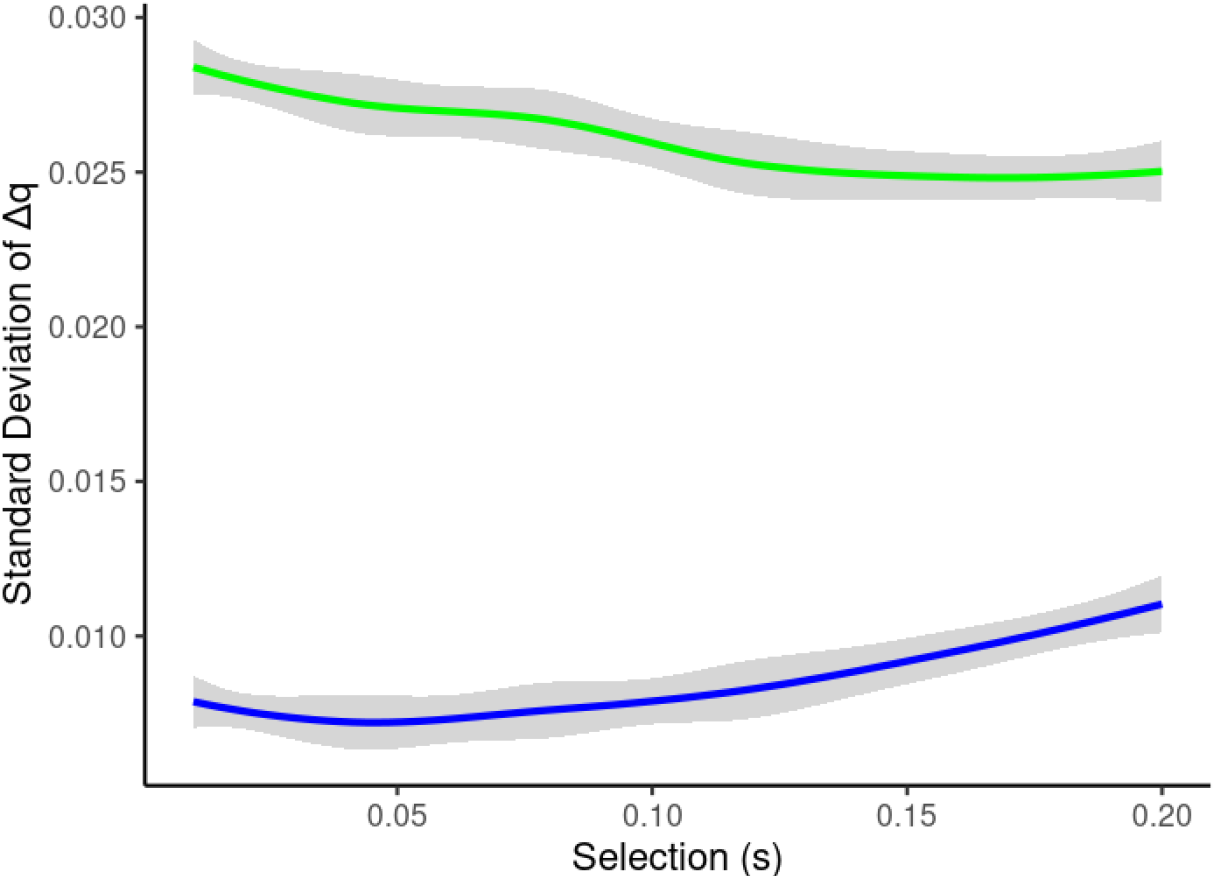
Standard deviation of *Δq* for beneficial mutations going to fixation from an initial frequency of *1/2N* across selection coefficients from *s = 0*.*01* to *0*.*20* for *N = 100* (green line) and *N = 1000* (blue line). The line represents the loess-smoothed means across 100 replications for each *N*, and gray shading indicates the 95% confidence interval.

**Fig. S4.**
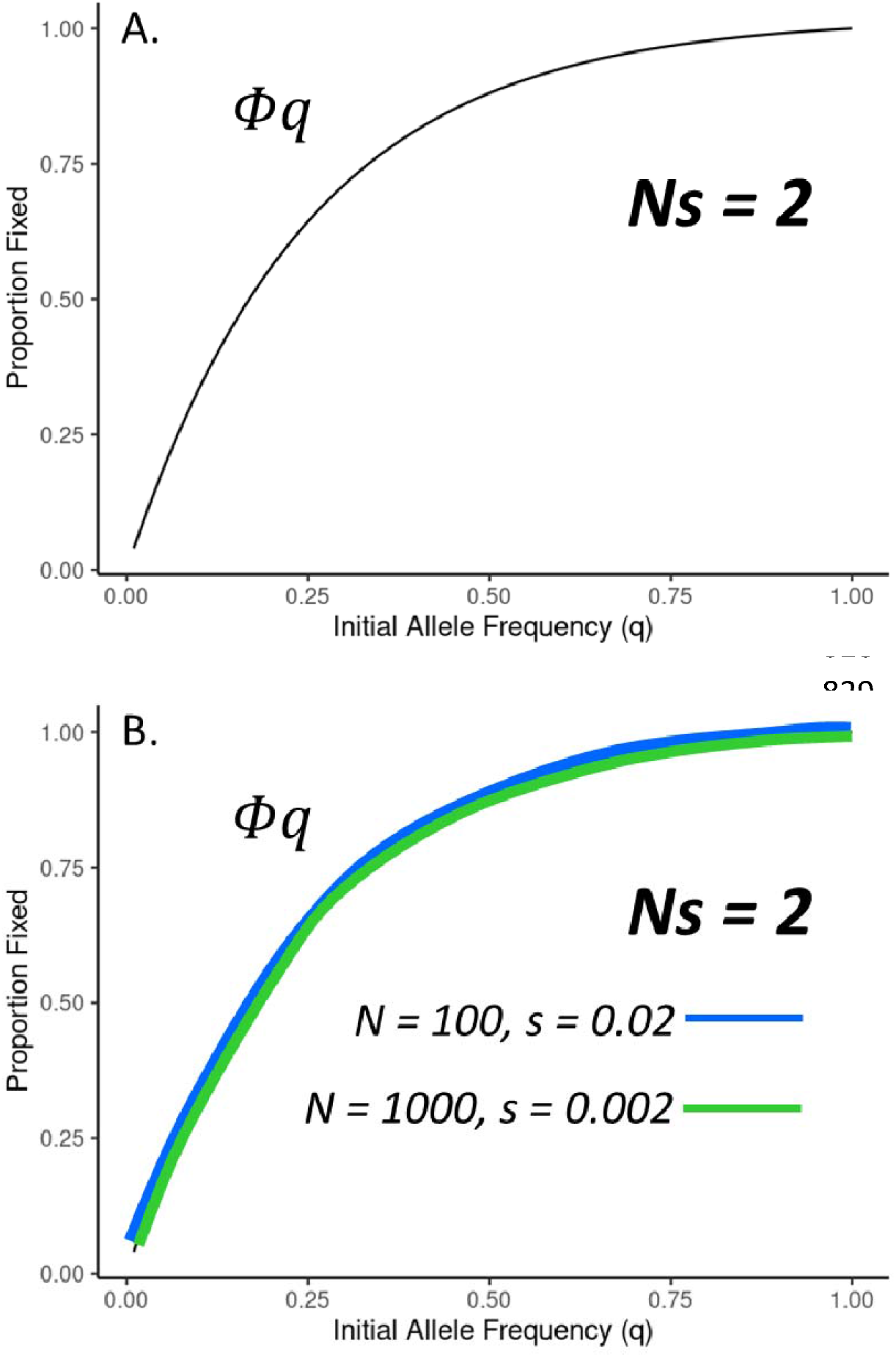
The diffusion approximation for the probability of fixation () in response to initial allele frequency with *Ns = 2 (A)*, and the proportion of fixed alleles across 100 loci after 1000 generations for population sizes of *N = 100* and 1000 with *Ns = 2* (B; the relationship for is obscured). Lines represent the loess-smoothed relationship of means for 20 replications per population size.

