## Supplementary material for "Testing Wright’s Intermediate Population Size Hypothesis – When Genetic Drift is a Good Thing": Scripts

Appendix 1. Python SimuPOP scripts.

A. **Track delta-q over generations for beneficial mutations**

import statistics

import numpy as np

reps = 50	#reps per N and s

repetitions = str(reps)

import simuPOP as sim

selection = [1, 4, 8, 12, 16, 20]

popsizes = [4, 8, 10, 20, 30, 40, 60, 80, 100, 120, 160, 200, 260, 320, 400, 500, 750, 1000, 2000, 3000]

for sel in range(0, len(selection)): #for each value of s

s = int(selection[sel])/100	#calculate s = 0.01 to 0.2

sel = str(s)

outfile = 'deltaq_by_N_and_s_h5_s=' + sel + 'reps=' + repetitions + '.txt' #define ouput file name

f = open(outfile, 'w')

f.write("	popsize	q	fit11	fit10	fit00	gens	mean	stdev \n") #column labels

f.close

for pop in range(0, len(popsizes)): #for each N

N = int(popsizes[pop])

initfreq = 1/(2*popsize) #initial frequency = 1/2N

fit00 = 1 - s	#normal allele fitness

fit10 = 1 - (s/2)	#heterozygote fitness (codominance h = 0.5 = 1 - s/2)

k = 0	#initiate reps

ntry = 0	#keep track of number of mutations lost

while k <= reps: #keep generating mutations until k=reps fix

ntry += 1

simu = sim.Simulator(sim.Population(N, loci=1), rep=1) #(N=population size. rep= number of populations
 simu.evolve(

initOps=[

sim.InitSex(maleProp=0.5), #equal sex ratio

sim.InitGenotype(freq=[(1 - initfreq), initfreq]), #set allele frequencies

sim.PyExec('qfreq = []')	#initiate list of mutation allele frequencies

],

matingScheme = sim.RandomMating(ops=[	#random mating

sim.MendelianGenoTransmitter(),

sim.MapSelector(loci=0, fitness={(0,0):fit00, (0,1):fit10, (1,1):fit11}), #fitness of each genotype

]),

postOps=[

sim.InitSex(maleProp=0.5), #equal sex ratio

sim.Stat(alleleFreq=0), #Generate statistics for locus 0

sim.PyExec('qfreq.append(alleleFreq[0][1])'), #list of allele freqs over generations

sim.TerminateIf('len(alleleFreq[0]) == 1') #terminate if the mutation is fixed or lost

],

)

qfreq1 = (simu.dvars(0).qfreq) #extract list of allele frequencies

ngens = len(qfreq1)	#get number of generations to fixation

if 1.0 in qfreq1: #only for mutations that fixed do this

k += 1	#increment k

deltaq = []	#initiate deltaq list

i = 1	#initiate generation counter i

while i < ngens: #calcuate delta-q

deltaq.append((qfreq1[(i)] - qfreq1[i-1])) #add delta-q to list

i += 1	#increment i

qmean = np.mean(deltaq) #calculate mean and standard deviation for fixed mutations

stdev = np.std(deltaq)

stqfreq1 = str(qfreq1)

f = open(outfile, 'a') #print results for fixed mutations

f.write(f" %10d %10.4f %10.4f %10.4f %10.4f %10d %10.4f %10.4f " % (N, initfreq, fit11, fit10, fit00, ngens, qmean, stdev))

f.write(" \n ")

print(N, s, k)	#keep track of progress by print to screen

B. **#Track q over generations with beneficial mutation initial frequency = 1/2N and Ns = 2**

#Import libraries and define variables

import statistics

import numpy as np

maxN = 1000

reps = 10000	#reps per N

outfile = "deltaq100reps_Ns2_Ns.txt"

f = open(outfile, 'w')

f.write("	popsize	q	fit11	fit10	fit00	gens	mean	stdev \n")

f.close()

import simuPOP as sim

popsizes = [10, 20, 30, 40, 60, 80, 100, 120, 160, 200, 260, 320, 400, 500, 750, 1000, 2000, 3000]

for pop in range(0, len(popsizes)):

popsize = int(popsizes[pop])

initfreq = 1/(2*popsize)

fit00 = 2/popsize

fit10 = 0.5*fit00	#dominance h = 0.5

k = 0

while k <= 10

simu = sim.Simulator(sim.Population(popsize, loci=1), rep=1) #(N=population size. rep= number of populations
 simu.evolve(

initOps=[

sim.InitSex(maleProp=0.5), #Make sure male/female ratios allow for offspring

sim.InitGenotype(freq=[(1 - initfreq), initfreq]), #set allele frequency

],

matingScheme = sim.RandomMating(ops=[

sim.MendelianGenoTransmitter(),

sim.MapSelector(loci=0, fitness={(0,0):fit00, (0,1):fit10, (1,1):fit11}), #fitness of each genotype [0, 0]

]),

postOps=[

sim.InitSex(maleProp=0.5), #Make sure male/female ratios allow for offspring

sim.Stat(alleleFreq=0), #Generate statistics for locus 0

#sim.PyExec('qfreq.append(alleleFreq[0][1])'),

sim.TerminateIf('len(alleleFreq[0]) == 1')

],

)

qfreq1 = (simu.dvars(0).qfreq) #extract list of q

ngens = len(qfreq1) #get number of generations to fixation

#select only runs where the mutation went to fixation

if 1.0 in qfreq1:

k += 1

deltaq = []	#initiate deltaq list

i = 1

while i < ngens: #calcuate delta-q

deltaq.append((qfreq1[(i)] - qfreq1[i-1])+1)

i += 1

qmean = np.mean(deltaq)

stdev = np.std(deltaq)

stqfreq1 = str(qfreq1)

f = open(outfile, 'a')

f.write(" %10d %10.4f %10.4f %10.4f %10.4f %10d %10.4f %10.4f " % (popsize, initfreq, fit11, fit10, fit00, ngens, qmean, stdev))

f.write(stqfreq1)

f.write(" \n ")

C. **#deleterious mutation accumulation with selection and drift**

#define variables, import libraries, create output file

import numpy as np

popsize = 0

initfreq = 0

maxN = 5000	#maximum population size

Npops = 1000	#number of populations (= number of loci)

reps = 100	#number of repetitions for each pop size

maxgen = 1000	#number of generations to run per rep

outfile = "deleterious_mutations(e-6).txt"

f = open(outfile, 'w')

f.write("	N	p	maxgen	Nloci	w(00) w(01) w(11) gens	freq1	freq0	fixed1	lost1 \n")

import simuPOP as sim

#generate a range of population sizes

while popsize < maxN:

if popsize < 10:

popsize +=2

elif popsize >= 10 and popsize < 30:

popsize +=5

elif popsize >= 30 and popsize < 80:

popsize +=10

elif popsize >= 80 and popsize < 200:

popsize +=20

elif popsize >= 200 and popsize < 500:

popsize +=50

elif popsize >= 500 and popsize < 1000:

popsize +=100

elif popsize >= 1000 and popsize < 10000:

popsize +=1000

#for each N replicate reps times

for x in range(reps):

s = np.random.poisson(2)/25 #random s from a Poisson distribution

h = np.random.uniform(0,0.2) #random h from a flat distribution

fit10 = 1-(h*s)

fit11 = 1-s

simu = sim.Simulator(sim.Population(popsize, loci=1), rep=Npops) #popsize=N, 1 locus per individual

simu.evolve(

initOps=[

sim.PyExec("f = 0"),

sim.InitSex(maleProp=0.5), #Make sure proportions allow for offspring

sim.InitGenotype(freq=[(1 - initfreq), initfreq]) #set allele frequency

],

preOps=sim.MatrixMutator(rate = [

[0, 1e-3],

[0, 0]

]),

matingScheme = sim.RandomMating(ops=[

sim.MendelianGenoTransmitter(),

sim.MapSelector(loci=0, fitness={(0,0):fit00, (0,1):fit10, (1,1):fit11}), #fitness of each genotype [0, 0]

]),

postOps=[

sim.InitSex(maleProp=0.5), #Make sure proportions allow for offspring

sim.Stat(alleleFreq=0), #Generate statistics for locus 0

#sim.TerminateIf('alleleFreq[0] == 1')

],

gen=maxgen

)

#extract information from simu.dvars and place in lists

allelefreq0 = [simu.dvars(x).alleleFreq[0][0] for x in range(Npops)] #locus 0 allele 0

allelefreq1 = [simu.dvars(x).alleleFreq[0][1] for x in range(Npops)] #locus 0 allele 1

popnum = [simu.dvars(x).rep for x in range(Npops)] #number of reps

popgen = [simu.dvars(x).gen for x in range(Npops)] #number of generations

meangen = sum(popgen)/len(popgen)

numlost0 = sum(map(lambda x : x == 0, allelefreq0))

numfixed0 = sum(map(lambda x : x == 1, allelefreq0))

numlost1 = sum(map(lambda x : x == 0, allelefreq1))

numfixed1 = sum(map(lambda x : x == 1, allelefreq1))

freq1 = sum(allelefreq1)/len(allelefreq1)

freq0 = sum(allelefreq0)/len(allelefreq0)

f = open(outfile, 'a')

f.write("%6d %10.4f %10d %10d %10.4f %10.4f %10.4f %10.4f %10.4f %10.4f %10d %10d \n" %

(popsize, initfreq, maxgen, Npops, fit00, fit10, fit11, meangen, freq1, freq0, numfixed1, numlost1))

f.close()

D. **#beneficial mutation with random selection (Poisson) and drift (uniform) initial q = 1/2N (no mutations)**

#Import libraries and define variables

import numpy as np

popsize = 0

initfreq = 0

maxN = 4000	#maximum population size

Npops = 1000	#number of populations

reps = 100	#number of repetitions for each pop size

maxgen = 1000	#number of generations to run per rep

outfile = "beneficial_mutations(e-6).txt"

f = open(outfile, 'w')

f.write("	N	p	maxgen	Nloci	w(00) w(01) w(11) gens	freq1	freq0	fixed1	lost1 \n")

import simuPOP as sim

#generate a range of population sizes

while popsize < maxN:

if popsize < 10:

popsize +=2

elif popsize >= 10 and popsize < 30:

popsize +=5

elif popsize >= 30 and popsize < 80:

popsize +=10

elif popsize >= 80 and popsize < 200:

popsize +=20

elif popsize >= 200 and popsize < 500:

popsize +=50

elif popsize >= 500 and popsize < 1000:

popsize +=100

elif popsize >= 1000 and popsize < 10000:

popsize +=1000

#for each N replicate reps times

for x in range(reps):

s = np.random.poisson(2)/25 #random s from a Poisson distribution

h = np.random.uniform(0,0.2) #random h from a flat distribution

fit10 = 1-(h*s)

fit00 = 1-s

initfreq = 1/(2*popsize)	#set initial mutation frequency

simu = sim.Simulator(sim.Population(popsize, loci=1), rep=Npops) #(N=population size. rep= number of populations

simu.evolve(

initOps=[

sim.PyExec("f = 0"),

sim.InitSex(maleProp=0.5), #Make sure male/female ratios allow for offspring

sim.InitGenotype(freq=[(1 - initfreq), initfreq]) #set allele frequency

],

matingScheme = sim.RandomMating(ops=[

sim.MendelianGenoTransmitter(),

sim.MapSelector(loci=0, fitness={(0,0):fit00, (0,1):fit10, (1,1):fit11}), #fitness of each genotype [0, 0]

]),

postOps=[

sim.InitSex(maleProp=0.5), #Make sure male/female ratios allow for offspring

sim.Stat(alleleFreq=0), #Generate statistics for locus 0

sim.TerminateIf('alleleFreq[0] == 1')

],

gen=maxgen

)

#extract information from simu.dvars and place in lists

allelefreq0 = [simu.dvars(x).alleleFreq[0][0] for x in range(Npops)]

allelefreq1 = [simu.dvars(x).alleleFreq[0][1] for x in range(Npops)]

popnum = [simu.dvars(x).rep for x in range(Npops)]

popgen = [simu.dvars(x).gen for x in range(Npops)]

meangen = sum(popgen)/len(popgen)

numlost0 = sum(map(lambda x : x == 0, allelefreq0))

numfixed0 = sum(map(lambda x : x == 1, allelefreq0))

numlost1 = sum(map(lambda x : x == 0, allelefreq1))

numfixed1 = sum(map(lambda x : x == 1, allelefreq1))

freq1 = sum(allelefreq1)/len(allelefreq1)

freq0 = sum(allelefreq0)/len(allelefreq0)

f = open(outfile, 'a')

f.write("%6d %10.4f %10d %10d %10.4f %10.4f %10.4f %10.4f %10.4f %10.4f %10d %10d \n" %

(popsize, initfreq, maxgen, Npops, fit00, fit10, fit11, meangen, freq1, freq0, numfixed1, numlost1))

f.close()

E. **#beneficial mutation with selection and drift for fixation flux**

\# define libraries and set variables

import numpy as np

Npops = 1000	#number of populations = independent loci (one locus per population)

reps = 10	#number of replications for each pop size

initfreq = 0	#1/(2*popsize)

maxgen = 4000	#number of generations to run per rep

outfile = "beneficial_mutation_Ns2_N4k.txt"

f = open(outfile, 'w')

f.write("	N	p	maxgen	Nloci	w(00) w(01) w(11) gens	freq1	freq0	fixed1	lost1 \n")

import simuPOP as sim

popsizes = [4, 6, 8, 10, 20, 30, 40, 60, 80, 100, 120, 160, 200, 260, 320, 400, 500, 750, 1000, 2000, 3000]

#for each N defined in list popsizes

for pop in range(0, len(popsizes)):

popsize = int(popsizes[pop])

for x in range(reps):

s = 2/popsize

h = 0.5	#Dominance additive h=0.5

fit10 = 1-(h*s)

fit00 = 1-s

simu = sim.Simulator(sim.Population(popsize, loci=1), rep=Npops) #(N=population size. rep= number of populations

simu.evolve(

initOps=[

sim.PyExec("f = 0"),

sim.InitSex(maleProp=0.5), #Make sure female/male ratios allow for offspring

sim.InitGenotype(freq=[(1 - initfreq), initfreq]) #set allele frequency

],

preOps=sim.MatrixMutator(rate = [	#mutation rate per generation

[0, 1e-4],

[0, 0]

]),

matingScheme = sim.RandomMating(ops=[

sim.MendelianGenoTransmitter(),

sim.MapSelector(loci=0, fitness={(0,0):fit00, (0,1):fit10, (1,1):fit11}), #fitness of each genotype [0, 0]

]),

postOps=[

sim.InitSex(maleProp=0.5), #Make sure female/male ratios allow for offspring

sim.Stat(alleleFreq=0), #Generate statistics for locus 0

#sim.TerminateIf('len(alleleFreq[0]) == 1')

],

gen = maxgen

)

#extract information from simu.dvars and place in lists

allelefreq0 = [simu.dvars(x).alleleFreq[0][0] for x in range(Npops)]

allelefreq1 = [simu.dvars(x).alleleFreq[0][1] for x in range(Npops)]

popnum = [simu.dvars(x).rep for x in range(Npops)]

popgen = [simu.dvars(x).gen for x in range(Npops)]

meangen = sum(popgen)/len(popgen)

numlost0 = sum(map(lambda x : x == 0, allelefreq0))

numfixed0 = sum(map(lambda x : x == 1, allelefreq0))

numlost1 = sum(map(lambda x : x == 0, allelefreq1))

numfixed1 = sum(map(lambda x : x == 1, allelefreq1))

freq1 = sum(allelefreq1)/len(allelefreq1)

freq0 = sum(allelefreq0)/len(allelefreq0)

f = open(outfile, 'a')

f.write("%6d %10.4f %10d %10d %10.4f %10.4f %10.4f %10.4f %10.4f %10.4f %10d %10d \n" %

(popsize, initfreq, maxgen, Npops, fit00, fit10, fit11, meangen, freq1, freq0, numfixed1, numlost1))

f.close()
